# Beyond Structure: Protein and Solution Dynamics Shape Ligand-Binding Thermodynamics

**DOI:** 10.64898/2026.09.29.755381

**Authors:** Kacie A. Evans, Morgan Powers, Robert L. Rider, Carter Lantz, Arthur Laganowsky, Hays S. Rye, David H. Russell

## Abstract

Variable-temperature electrospray ionization native mass spectrometry (vT-ESI nMS) provides a means to determine how temperature-dependent changes in protein and solution dynamics influence individual ligand-binding reactions within multiligand protein complexes. Here, the GroEL single-ring mutant (SR1), which binds up to seven nucleotides, serves as a model system due to its sensitivity to solution conditions. Native MS resolves individual ligand-bound populations, while vT-ESI extends these measurements across temperature (3-43 °C), enabling determination of binding affinities and the associated Gibbs free energy, enthalpy, entropy, heat capacity changes (ΔC_p_), and enthalpy-entropy compensation (EEC) for each binding reaction. Temperature-dependent changes in van’t Hoff curvature produce distinct ΔC_p_ profiles, indicating changes in the dynamics governing ligand binding that may reflect contributions from protein conformational and protonation microstates. Temperature-dependent shifts in average charge state, consistent with changes in solvent-accessible surface area, indicate corresponding changes in protein conformation. Binding is predominantly enthalpy-driven below ∼23 °C, with increasing entropic contributions at higher temperatures, while the seventh ADP binding reaction exhibits a distinct EEC profile. Comparison of the seventh ADP binding reaction in H_2_O and D_2_O reveals pronounced differences in van’t Hoff curvature, ΔC_p_, and EEC, demonstrating that changes in the hydration environment alter the thermodynamic response. Collectively, these measurements show that temperature-dependent changes in protein and solution dynamics alter the distributions and thermodynamics of individual ligand-bound states and demonstrate the utility of vT-ESI nMS for resolving these effects in complex multiligand systems.

## Introduction

Protein-ligand interactions are sensitive to solution conditions, including temperature,^1-3^ pH,^4^ buffer composition,^5^ osmolytes,^6^ hydration,^7^ and pressure.^8^ These environmental factors influence both ligand binding affinity and the molecular mechanisms underlying protein-ligand recognition. Thermodynamic analysis provides a quantitative framework for characterizing these effects;^9^ however, conventional ensemble-averaged techniques, such as isothermal titration calorimetry (ITC)^10^ and nuclear magnetic resonance (NMR) spectroscopy,^11^ can be limited in resolving heterogeneous or temperature-dependent behavior among individual ligand-bound and unbound states. These limitations highlight the need for experimental approaches capable of resolving individual ligand-bound states and their temperature-dependent thermodynamics within complex multiligand systems.^12, 13^ Variable-temperature electrospray ionization (vT-ESI) native mass spectrometry (nMS) provides this capability by resolving individual ligand-bound populations, enabling sequential binding reactions to be characterized one ligand at a time.^3, 14, 15^ The importance of temperature as an experimental variable has long been recognized in protein crystallization, where careful temperature control is required because protein behavior is strongly temperature-dependent.^16^ Rather than treating temperature solely as a parameter to control, variable temperature approaches can exploit this sensitivity to investigate how temperature-dependent changes in protein and solution dynamics influence ligand binding, particularly in complex systems with multiple binding events.^3, 5, 14^ GroEL, a well-characterized molecular chaperonin involved in protein folding, provides a model multiligand system for investigating these effects.^17-19^ The GroEL single ring mutant (SR1) simplifies analysis of nucleotide binding by eliminating inter-ring cooperativity, while the use of ADP avoids complications associated with ATP hydrolysis.^19, 20^ Together, these features establish SR1-ADP as a well-defined model for investigating how solution conditions influence the thermodynamics of individual nucleotide-binding reactions within a multiligand system.

Protein-ligand binding occurs within a dynamic solvent environment in which hydration contributes to protein conformation, stability, and dynamics.^21-25^ Hydration influences conformational ensembles, stability, and ligand binding through reorganization of hydrogen-bonding networks and interactions with both polar and nonpolar regions of the protein.^26-29^ Temperature is also a key factor determining these effects, as it alters protein-water dynamics, which have been described in terms of distinct regimes of “cold” (more structured, less dynamic) and “hot” (less structured, more dynamic) water.^30-32^ Temperature-dependent changes in water structure are closely coupled to protein behavior, as increased interaction with hydrophobic regions at lower temperatures and reduced solvent organization at higher temperatures can shift protein dynamics and stability.^25, 33, 34^ As a result, temperature-dependent changes in hydration can influence protein-ligand binding thermodynamics through coupled changes in solvent organization and protein dynamics. Perturbing the hydration environment provides a means to examine its contribution to protein dynamics and ligand binding. Substitution of H_2_O with D_2_O alters hydrogen-bond strength and solvent dynamics, resulting in stronger and more persistent hydrogen-bonding networks.^35-38^ These differences make D_2_O a useful probe for investigating how hydration and solvent dynamics influence protein-ligand binding.^7, 39^

Previous vT-ESI nMS studies demonstrated that GroEL-nucleotide binding is sensitive to both temperature and ESI buffer composition, which alters nucleotide-binding affinities and cooperative binding behavior.^3, 5, 19^ While these studies establish that GroEL-nucleotide binding is strongly influenced by changes in the solution environment, how the effects of temperature alter the thermodynamics remains unresolved. Here, vT-ESI nMS was used to investigate how temperature-dependent changes in hydration and protein dynamics influence individual SR1-ADP binding reactions. ADP binding was monitored across temperature in H_2_O and D_2_O, enabling comparison of temperature-dependent binding thermodynamics under different hydration environments. Temperature-dependent measurements of average charge state (Z_avg_), van’t Hoff behavior, changes in heat capacity (ΔC_p_), and enthalpy-entropy compensation (EEC) were integrated to evaluate changes in protein and solution dynamics associated with SR1-ADP binding. Comparison of these behaviors in H_2_O and D_2_O further provided a means to evaluate how differences in hydration and solvent dynamics influence the binding thermodynamics.^7, 39-41^ Collectively, these measurements establish a framework for understanding how temperature-dependent protein and solution dynamics shape individual ligand-binding reactions within a multiligand system.

## Results

SR1-ADP binding was examined by vT-ESI nMS in H_2_O and D_2_O to determine how temperature and hydration influence individual nucleotide-binding reactions. All experiments were conducted in 200 mM ammonium acetate (AmAc), as previous studies showed that ethylenediamine diacetate (EDDA) and triethylammonium acetate (TEAA) inhibit cooperative nucleotide binding.^5, 19^ These experiments were performed by measuring Z_avg_ and ADP-bound populations at 2 °C temperature intervals, enabling evaluation of temperature-dependent conformational changes and determination of ADP binding affinities. Temperature-dependent binding affinities were used to construct van’t Hoff plots, from which ΔC_p_ and the enthalpic and entropic contributions to binding were determined. Together, these measurements revealed a common temperature-dependent change in SR1-ADP binding behavior near ∼23 °C, providing a basis for examining the protein and solution dynamics associated with this behavior.

Temperature-dependent SR1 dynamics were evaluated by monitoring shifts in the Z_avg,_ which report changes in solvent accessible surface area (SASA) for SR1 and SR1(ADP)_*n*_ (*n* = 1-7) complexes.^42, 43^ Temperature-dependent Z_avg_ plots for SR1(ADP)_3,5,7_ in H_2_O and D_2_O over 3-43 °C are shown in **Figure 1A,E**; temperatures above 45 °C were excluded as the GroEL/SR1 complex begins to dissociate into monomers.^5, 39^ The complete data sets for all SR1(ADP)_*n*_ complexes are provided in Figure S1. Across all temperatures, Z_avg_ decreased monotonically with increasing ADP binding (SR1(ADP)_3_ > SR1(ADP)_5_ > SR1(ADP)_7_) in both solvents. From 4 °C to ∼23 °C, Z_avg_ decreased slightly with increasing temperature across all SR1 and SR1(ADP)_*n*_ complexes, in both H_2_O and D_2_O, as shown in Figure S2. Above ∼23 °C, this trend reversed, and Z_avg_ increased by ∼2 charge states in both H_2_O and D_2_O. Differences in the separation of the mean Z_avg_ values among ADP-bound states are also apparent between H_2_O and D_2_O, although the associated error ranges overlap for some measurements. This reversal in the temperature dependence of Z_avg_ near ∼23 °C has been observed previously and is consistent with a change in the temperature-dependent conformational dynamics that may be influenced by reorganization of the surrounding hydration network.^5, 39^ Together, these results identify a change in the SR1 conformational ensemble near ∼23 °C, which may also involve changes in the distribution of protein protonation microstates.^44^

**Figure 1.**
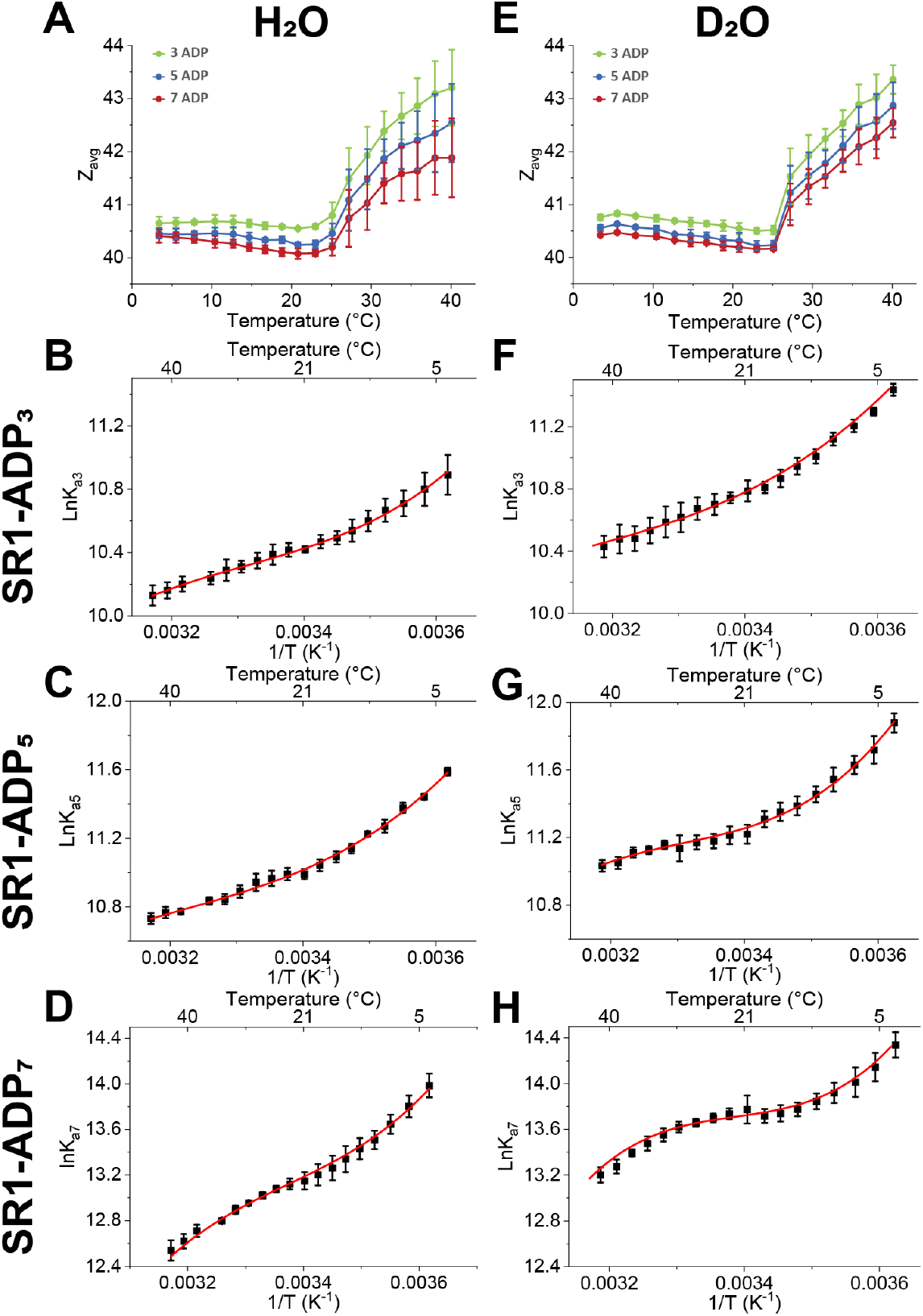
Effects of solution temperature contained in the ESI emitter on the Z_avg_ for SR1(ADP)_3,5,7_ in (**A**) H_2_O and (**E**) D_2_O with 20 µM ADP. The van’t Hoff plots are shown in (**B-D**) H_2_O and (**F-H**) D_2_O for SR1(ADP)_3,5,7_ complexes. Above ∼25 °C, Z_avg_ exhibits a greater temperature-dependent increase in H_2_O than in D_2_O, with greater variability observed in H_2_O. All values are generated from triplicate data sets, and error bars represent standard deviation (*n* = 3).

Temperature-dependent binding constants (K_a_) for SR1(ADP)_3,5,7_ in H_2_O and D_2_O were used to generate van’t Hoff plots shown in **Figure 1**; complete data sets for all SR1(ADP)_*n*_ complexes are provided in Figures S2 and S3. Nonlinear van’t Hoff behavior indicates a temperature-dependent ΔC_p_ likely arising from water reorganization or changes in protein conformation.^45^ In contrast to nonlinear van’t Hoff plots characterized by two distinct slopes, these data exhibit more complex curvature that was described using polynomial fits, with a third-order polynomial used for binding reactions exhibiting more complex curvature. A change in van’t Hoff curvature is apparent near 0.0034 K^−^ (∼23 °C) across the ADP binding reactions in both H_2_O and D_2_O, with the effect most pronounced for the 7^th^ ADP binding reaction (**Figure 1C,F**), particularly in D_2_O. For the 7^th^ ADP binding, the van’t Hoff plot transitions from convex curvature at lower temperatures to concave curvature at higher temperatures, indicating a pronounced temperature dependence of ΔC_p_. Together, these results identify a change in SR1–ADP binding thermodynamics near ∼23 °C that coincides with the change in temperature-dependent Z_avg_ behavior observed over the same temperature range.

Temperature-dependent ΔC_p_ values were determined from the curvature of the van’t Hoff plots and are shown in **Figure 2** for SR1(ADP)_5,6,7_ complexes in H_2_O and D_2_O. ΔC_p_ vs. T plots for all SR1(ADP)_*n*_ complexes are provided in Figures S4 and S5. For most, but not all, ADP bindings, ΔC_p_ was positive at lower temperatures and became negative at higher temperatures in both solvents, with convex and concave curvature corresponding to positive and negative ΔC_p_, respectively.

**Figure 2.**
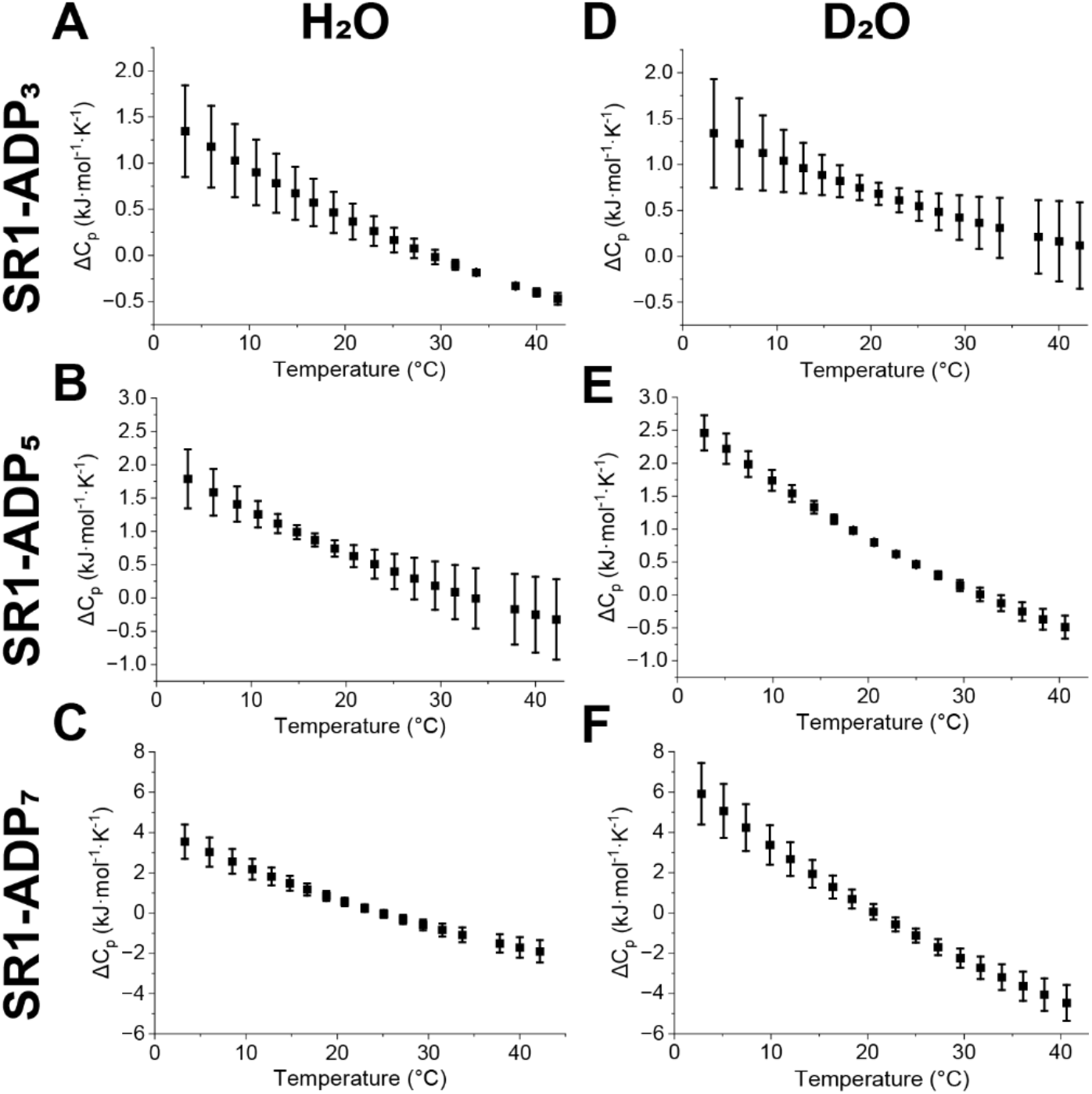
Plots of ΔC_p_ in relation to temperature are shown in (**A-C**) H_2_O and (**D-F**) D_2_O for SR1(ADP)_3,5,7_ complexes. All values are generated from triplicated data sets, and error bars represent standard deviation (*n* = 3).

Changes in ΔC_p_ reflect temperature-dependent changes in the ensemble of interactions contributing to binding, which can include protein conformational changes, hydration and hydrogen-bond network reorganization, and other weak interactions within the protein-ligand-solution system.^46, 47^ The magnitude of ΔC_p_ increases as ADP binds to SR1, indicating that later ADP bindings are associated with increasingly pronounced temperature-dependent changes in the interactions and dynamics contributing to binding. The magnitude of ΔC_p_ was also consistently greater in D_2_O than in H_2_O across the measured temperature range. Larger uncertainties in ΔC_p_ at the temperature extremes likely reflect increased sensitivity of the thermodynamic response in regions where hydration-mediated interactions and protein conformational dynamics undergo the greatest temperature-dependent changes. The temperature-dependent shift in ΔC_p_ from positive to negative values provides a quantitative basis for the transition observed in the van’t Hoff analysis, indicating continual reorganization of protein-solvent and protein-ligand interactions as the balance between structured hydration and conformational flexibility changes with temperature. Importantly, the temperature-dependent ΔC_p_ behavior coincides with the change in Z_avg_ near ∼23 °C, linking changes in SR1 conformational dynamics with changes in the thermodynamic response of ADP binding. Overall, these results indicate that temperature-dependent changes in hydration and protein dynamics influence SR1-ADP binding thermodynamics as additional ADP binds to SR1 and are amplified in D_2_O solutions.

Temperature-dependent enthalpy (ΔH), entropy (-TΔS), and free energy (ΔG) profiles derived from van’t Hoff analysis for SR1(ADP)_3,5,7_ complexes are shown in **Figure 3**. ΔH, -TΔS, and ΔG vs. T plots for all SR1(ADP)_*n*_ complexes are shown in Figures S6 and S7. At low temperatures, ligand binding is predominantly enthalpy-driven for all ADP bindings in both H_2_O and D_2_O. As the temperature approaches ∼23 °C, corresponding to the inflection point observed in the van’t Hoff plots, the enthalpic and entropic contributions approach similar magnitudes. For most ADP bindings, binding becomes increasingly entropy-favored at higher temperatures. Most notably, the 7^th^ ADP binding exhibits distinct behavior relative to the other ADP bindings. In H_2_O, the 7^th^ binding does not fully transition to an entropy-dominated regime but instead reaches a balance between enthalpy and entropy before returning to enthalpy dominance at higher temperatures. In D_2_O, the 7^th^ binding displays two transition temperatures, with entropy being more favorable at intermediate temperatures before shifting back to enthalpy dominance at higher temperatures. These results indicate that the thermodynamic driving factors governing SR1-ADP binding change substantially across the temperature range and become increasingly complex for the final ADP binding event. Collectively, the temperature-dependent Z_avg_ trends, van’t Hoff analysis, ΔC_p_, and thermodynamic parameters reveal a transition near ∼23 °C in SR1-ADP binding consistent with coordinated changes in hydration and protein conformational dynamics.

**Figure 3.**
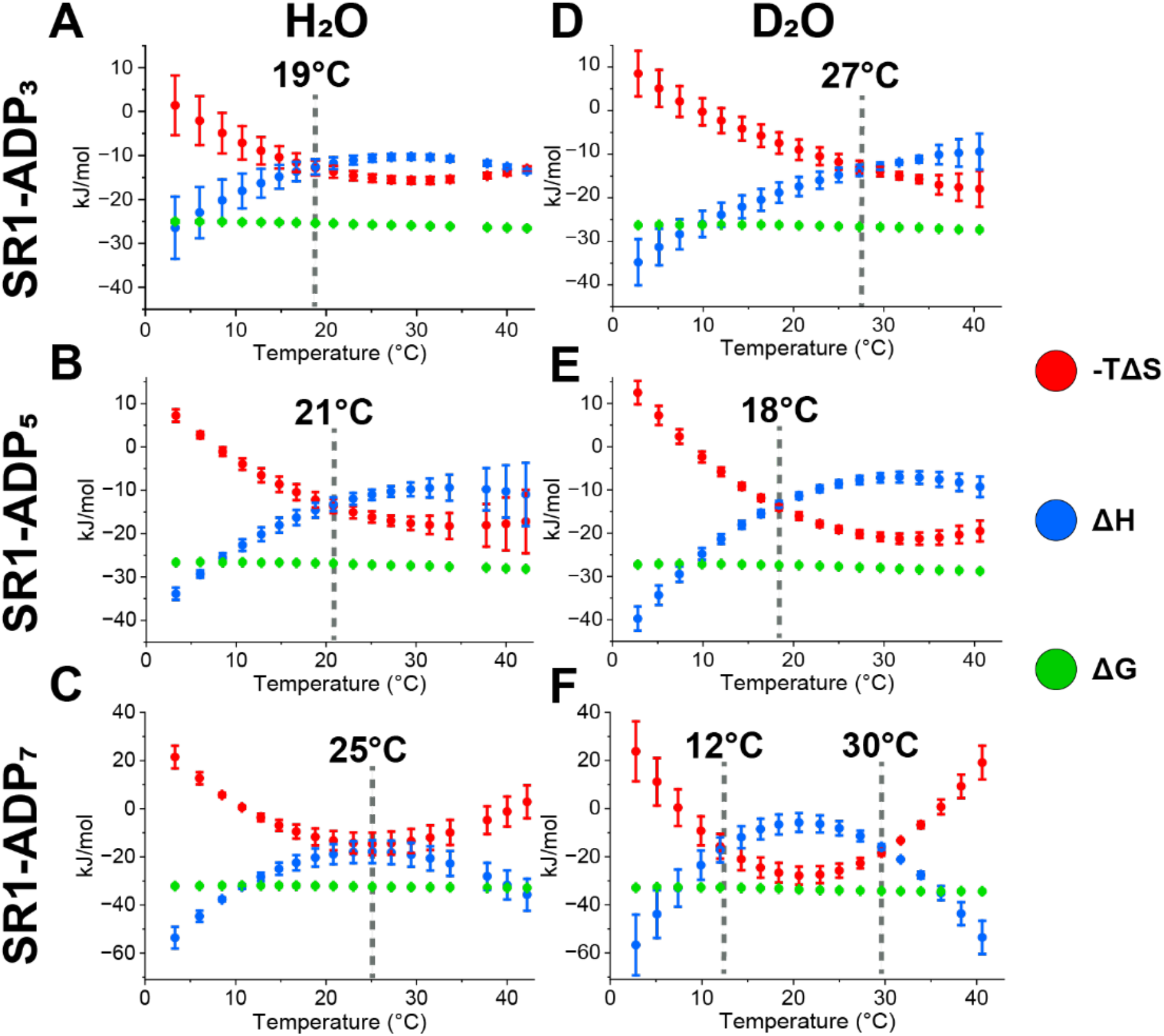
Plots of -TΔS, ΔH, and ΔG in relation to temperature are shown in (**A-C**) H_2_O and (**D-F**) D_2_O for SR1(ADP)_3,5,7_ complexes. Dashed gray lines indicate temperatures at transition points where -TΔS and ΔH crossover. All values are generated from triplicated data sets, and error bars represent standard deviation (*n* = 3).

## Discussion

Temperature-dependent measurements reveal that the coupled dynamics of the protein, ligand, and surrounding solution environment play a central role in modulating nucleotide binding to SR1. Together, temperature-dependent changes in Z_avg_, ΔC_p_, and EEC identify a distinct change in SR1-ADP binding behavior near ∼23 °C, reflecting a shift in the dynamics governing nucleotide binding. Comparison of H_2_O and D_2_O further shows that differences in hydration dynamics and solvent-mediated interactions alter these behaviors, emphasizing the role of the local solution environment in shaping binding thermodynamics. The corresponding shifts in enthalpic and entropic contributions suggest that temperature alters the balance among protein-solvent, ligand-solvent, and protein-ligand interactions.

Temperature-dependent thermodynamics shown in **Figure 3B-D** suggest that changes in hydration and protein dynamics modulate ADP binding to the SR1 complex. At lower temperatures, ADP binding is enhanced, particularly for the 7^th^ binding reaction, suggesting that structured hydration contributes to the interactions underlying cooperative nucleotide binding.^3, 5^ Positive ΔC_p_ values at lower temperatures (**Figure 2A-C**) together with the higher Z_avg_ observed at 4 °C relative to 23 °C (**Figure 1A**) are consistent with changes in hydration and slight expansion of the protein conformation within this low-temperature regime. These observations suggest that the low-temperature regime favors increased protein-solvent interactions and a more structured hydration environment surrounding the ADP-bound complex.^28, 46^ Binding in this regime is predominantly enthalpy-driven, consistent with reduced thermal motion and more persistent hydration networks that favor stabilizing protein-solvent and protein-ligand interactions.^28^ At higher temperatures, ΔC_p_ becomes negative while Z_avg_ increases (**Figure 2A-C** and **Figure 1A**), consistent with increased conformational flexibility and altered solvent interactions.^47-51^ The accompanying increase in the entropic contribution to binding (**Figure 3A,B)** is consistent with greater conformational sampling and reduced solvent organization, accompanied by redistribution of the surrounding hydration environment.^52^ The EEC crossovers observed at 19 °C and 21 °C for the intermediate ADP binding reactions (**Figure 3A,B**) mark temperatures where enthalpic stabilization becomes balanced by increasing entropic contributions from the coupled protein-ligand-solution system.^52^ These shifts are consistent with temperature-dependent reorganization of water, AmAc ions, and protein conformations that alters both the interactions and configurational freedom contributing to ADP binding. This behavior is consistent with two-state models of water, in which temperature-dependent reorganization between more structured and more dynamic hydrogen-bonding regimes alters hydration dynamics and biomolecular interactions.^29-31, 53-55^ Rather than occurring abruptly, this change extends over a narrow temperature range and resembles non-Arrhenius-like behavior arising from temperature-dependent changes in protein and solvent interactions.^56, 57^ The 2 °C temperature sampling used here enables these gradual changes to be resolved. Together, these observations indicate that the change in SR1-ADP binding near ∼23 °C reflects a gradual redistribution of the dynamics and interactions within the coupled protein-ligand-solution system.

The larger uncertainties in ΔC_p_ at the temperature extremes reflect increased sensitivity of the thermodynamic response to changes in the underlying molecular ensemble. Within this ensemble, changes in protein conformation alter microscopic protonation equilibria, shifting the distribution of protein protonation microstates. As described by Khaniya et al., coupling between conformational and protonation states produces microscopic states with distinct energetic contributions.^44^ This interpretation is also consistent with Cooper’s analysis of protein heat capacities, which demonstrated that large heat-capacity changes are not unique to proteins but can accompany transitions between molecular states with different degrees of order and accessible configurations.^47^ Thus, temperature-dependent changes in hydration, protein conformation, and protonation equilibria alter the ensemble contributing to the measured thermodynamic response, resulting in greater ΔC_p_ variability at the temperature extremes.

The temperature-dependent thermodynamic response becomes increasingly pronounced with ADP occupancy, with the 7^th^ ADP binding reaction exhibiting the greatest sensitivity (**Figure 3C**). As additional ADP binds, SR1 adopts a more extended conformation, increasing solvent accessibility and potentially strengthening the coupling between protein conformational dynamics and the surrounding solution environment. Structural studies have shown that ADP binding induces a partially extended conformation of GroEL that resembles the ATP-bound state, although to a lesser extent,^20^ while SR1 samples a more flexible conformational ensemble capable of adopting expanded states.^58^ The distinct thermodynamic behavior of the 7^th^ ADP binding reaction (**Figure 3C**) further reflects the cooperative nature of GroEL, in which ligand binding is coupled across subunits.^20, 58-61^ Unlike the earlier ADP bindings (**Figure 3A,B**), the final ADP binding does not exhibit a distinct crossover from enthalpy-to entropy-dominated binding. Instead, enthalpic and entropic contributions remain broadly balanced across ∼18-32 °C (**Figure 3C**), suggesting that protein, ligand, and surrounding solution interactions continue to reorganize over this temperature range rather than undergoing a single thermodynamic crossover. Increased solvent accessibility in the more extended SR1 ensemble may facilitate redistribution or exchange of water within solvent-accessible regions, increasing the sensitivity of the final binding reaction to temperature-dependent changes in hydration and protein dynamics. Similar hydration-coupled behavior has been proposed for rhodopsin activation, where conformational expansion facilitates water influx and contributes to entropic stabilization.^62-64^ At higher temperatures, the return toward enthalpy-dominated binding may reflect stabilization through cooperative protein-protein and protein-ligand interactions.^18, 65, 66^ These results indicate that the enhanced sensitivity of the 7^th^ ADP binding arises from increased solvent exposure and cooperative interactions across subunits at the final binding step, amplifying the impact of temperature-dependent changes in protein conformational dynamics and hydration.

To further evaluate the contribution of hydration to the temperature-dependent thermodynamics of SR1-ADP binding, experiments were performed in D_2_O. Enhanced cooperative ligand binding has been observed in D_2_O,^39^ indicating that solvent-mediated interactions play an important role in stabilizing the SR1(ADP) complex. The lower zero-point vibrational energy of deuterium results in slightly stronger and longer-lived O-D hydrogen bonds and altered solvent properties, including viscosity, density, and hydration dynamics.^36-38^ Previous studies have shown that the enhanced protein stability observed in D_2_O is primarily a solution-phase phenomenon, supporting the interpretation that the thermodynamic differences observed here arise predominantly from altered hydration rather than direct protein isotope effects.^7, 67^ The greater temperature dependence observed in D_2_O (**Figure 3D-F**) suggests that perturbation of these more persistent hydration networks produces greater van’t Hoff curvature and a more pronounced thermodynamic response. This difference is particularly evident for the seventh ADP binding reaction (**Figure 3C,F):** whereas H_2_O exhibits a broad temperature range over which enthalpic and entropic contributions remain nearly balanced, D_2_O exhibits two distinct EEC crossover temperatures. The longer-lived O-D hydrogen-bonding network can slow reorganization of water surrounding the protein and solution ions.^68^ One possible interpretation is that slower interconversion among hydration states reduces the averaging that occurs in H_2_O, allowing thermodynamic regimes that appear as a broad compensation region in H_2_O to become resolved as two distinct crossover temperatures in D_2_O.^7^ The buffer environment may further contribute to this behavior, as ions can alter hydrogen-bonding networks and water dynamics, consistent with previous observations that the temperature-dependent Z_avg_ transition occurs in AmAc but not in EDDA or TEAA.^5, 69^ The enhanced curvature in the van’t Hoff plots and larger ΔC_p_ responses observed in D_2_O indicate that altering the hydration environment amplifies the temperature-dependent thermodynamic response of SR1-ADP binding, emphasizing the role of hydration dynamics and solvent-mediated interactions in modulating ligand binding thermodynamics. Collectively, these results indicate that nucleotide binding emerges from the coupled dynamics of the protein, ligand, and surrounding solution environment, with changes in hydration and protein dynamics becoming increasingly important as cooperative binding progresses.

## Conclusions

The results presented here demonstrate that the SR1-ADP binding is strongly influenced by temperature through changes in the coupled dynamics of the protein, ligand, and surrounding solution environment. A consistent transition near ∼23 C is observed across temperature-dependent Z_avg_, van’t Hoff analysis, and ΔC_p_, indicating a shift in the balance of interactions governing ligand binding. This transition is consistent with models describing temperature-dependent changes in water structure and hydrogen-bonding networks, including two-state descriptions of hydration and studies highlighting the role of solvent dynamics in modulating protein behavior.^30, 53, 70^ As temperature increases, binding shifts from predominantly enthalpy-driven at lower temperatures to greater entropic contributions, reflecting continual reorganization of the protein-ligand-solution system. These observations suggest that structured hydration contributes more significantly to binding at lower temperatures, whereas increased thermal motion at higher temperatures alters both protein dynamics and solvent organization.^29, 32^ Comparison with D_2_O further highlights the importance of the surrounding solution environment, as stronger hydrogen-bonding networks alter the collective dynamics of hydration, buffer ions, and the protein, amplifying the temperature-dependent thermodynamic response and observed transition behavior.^37, 38^

The intrinsic temperature dependence of SR1 nucleotide binding suggests that sensitivity of ligand-binding thermodynamics to the solution environment may provide a means for chaperonins to adapt their energetic landscape across changing solution conditions.^3, 71, 72^ Collectively, these findings demonstrate that temperature-dependent ligand-binding thermodynamics emerge from the coupled dynamics of the protein and surrounding solution environment. More broadly, these results demonstrate the ability of vT-ESI native mass spectrometry to resolve temperature-dependent thermodynamics for individual ligand-binding reactions while simultaneously monitoring changes associated with protein and solution dynamics. This capability creates opportunities to integrate vT-ESI nMS with complementary solution-phase measurements, such as single-molecule Förster resonance energy transfer (smFRET), to directly relate ligand-binding thermodynamics to conformational states and dynamics. Recent integration of vT-ESI nMS and smFRET for GroEL demonstrates the potential of this combined approach to connect ligand binding with allosteric conformational dynamics.^73^ Together, these complementary approaches provide a framework for investigating how protein, ligand, and solution dynamics shape biomolecular interactions.

## Methods

### Sample Preparation

All chemicals, including AmAc, ADP, and 2M acetate (MgAc_2_) were purchased from Sigma-Aldrich (St. Louis, MO) and were dissolved in LC-MS grade deionized water. AmAc was dissolved directly in LC-MS grade deionized H_2_O or D_2_O to prepare 200 mM AmAc solutions in H_2_O and D_2_O, respectively. All AmAc buffer solutions (H_2_O and D_2_O) were adjusted to pH 7 using ammonium hydroxide. SR1 was overexpressed in E. coli as described previously.^74^ Aliquots of ADP solutions containing 1 mM MgAc_2_ were stored at -20 °C and freshly diluted with the pH-adjusted 200 mM AmAc buffer containing 1 mM MgAc_2_, then added to the protein prior to analysis. ADP concentration was measured by UV-Vis at 259 nm. Protein concentration was measured by UV-Vis at 280 nm. Fresh SR1 was diluted 3-fold and buffer exchanged into the corresponding buffer containing 1 mM MgAc_2_ using a Micro Bio spin P-6 gel column (Bio-Rad). Deuterated samples were incubated at 4 °C for 48 h to allow hydrogen-deuterium exchange to reach equilibrium prior to analysis, resulting in approximately 53-55% exchange of the total exchangeable hydrogens in the complex (see Figures S8 and S9).

### Variable-Temperature Native Mass Spectrometry Analysis

The temperature of the solution contained in the nano-ESI emitter was controlled by the home-built variable temperature device as described previously.^14^ The vT-ESI temperature suggests an error of ± 1.5 °C. Solution temperatures used for this study were 3 - 43 °C. Solution temperatures were measured using a calibrated T-Type thermocouple (Physitemp Clifton, NJ) paired to a thermocouple (National Instruments USB-TC01) positioned inside the borosilicate pulled glass capillary filled with 1 µM SR1 and 1 mM MgAc_2_ in 200 mM AmAc in H_2_O or D_2_O. The actual temperature in relation to the input temperature for H_2_O and D_2_O is shown in Figure S10. ADP solutions at various concentrations prepared in the same buffer as SR1 were titrated into SR1 and incubated at each temperature for 2 min. Then, mass spectra were collected on a Thermo Q Exactive UHMR (ultra-high mass range) hybrid quadrupole orbitrap mass spectrometer. The resolution setting was maintained at 12500 with 5 microscans for SR1-ADP binding experiments. The capillary temperature was set to 100 °C with in-source trapping set to −200 V, and the HCD energy was set to 220. Using these conditions, no gas-phase dissociation products were observed. The acquisition time for each spectrum was set to 1 min.

### Data Processing

UniDec was used to assign the charge states, mass, and abundance of each individual species detected in the mass spectra.^75^ Mass assignment information for all species is provided in Tables S1 and S2. Z_avg_ was calculated as the weighted average of all charge states for a mass species. The integrated signal intensities of each complex were used to fit a sequential binding model for solving dissociation constant (K_d_) values as previously described by Cong et al.,^76^ from which the apparent binding constants (or the equilibrium constant K_eq_) are obtained as the reciprocals. The intrinsic binding constants (K_a_) are obtained using the equations reported from our past studies.^5, 19, 39^ The K_a_ values were used for the nonlinear van’t Hoff analysis to determine the temperature-dependent thermodynamic parameters. The Gibbs free energy for ADP binding was calculated using eq. 1.

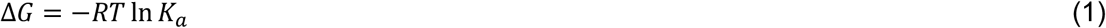

Where \**R* = 8.314 J·K^-1^·mol^-1^ and T is the absolute temperature in Kelvin. Van’t Hoff plots (*lnK*_*a*_ vs. *1/T*) were fit using a second or third order polynomial functions (OriginPro 2025, Simple Fit application). The polynomial order was selected based on improvement in the coefficient of determination (R^2^) and inspection of residuals to ensure no systematic deviations.

Temperature-dependent enthalpy values were obtained from the derivative form of the van’t Hoff equation (eq. 2).

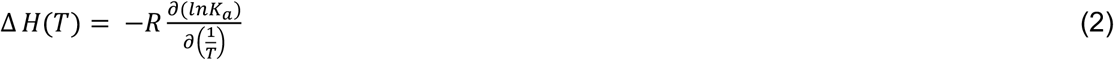

The corresponding ΔC_p_ was calculated using eq. 3.

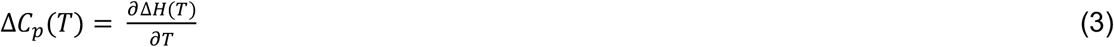

The temperature dependence of entropy is related to the ΔC_p_ according to 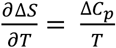, illustrating the relationship between ΔC_p_ and the temperature-dependent entropic contribution to binding.

Entropy values at a given temperature (*T*) were calculated from ΔH and ΔG using eq 4.

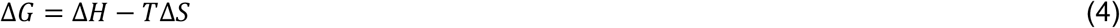

## Acknowledgement

Funding for this work was provided by the Robert A. Welch Foundation (Grant A-2106-20220331 to A.L. and Grant A-2162 to D.H.R.), National Institutes of Health (Grant RM1GM145416 to A.L., Grant R01GM134063-01 to H.R., and Grant RM1GM149374 to D.H.R.), Texas A&M University Division of Research Targeted Proposal Teams (TPT) funding program (H.R.), the Office of TAMU Vice-President for Research (D.H.R.), and the MDS Sciex Professor of Mass Spectrometry (D.H.R.).

## Supporting Information

Z_avg_ vs Temperature plots for SR1(ADP)_n_ complexes in H_2_O and D_2_O, van’t Hoff plots for SR1(ADP)_n_ complexes in H_2_O and D_2_O, ΔC_p_ vs temperature plots for SR1(ADP)_n_ complexes in H_2_O and D_2_O, Plots of ΔG, ΔH, and -TΔS vs temperature in H_2_O and D_2_O, Intact hydrogen deuterium exchange of SR1, mass spectra of SR1 in H_2_O and D_2_O after 48 h, actual vs input temperature plot for vT-ESI device for H_2_O and D_2_O, and mass assignment information of SR1(ADP)_n_ complexes in H_2_O and D_2_O.

## Author Information

### Authors

**Kacie A. Evans** - Department of Chemistry, Texas A&M University, College Station, Texas, 77843, United States

**Morgan Powers** - Department of Biochemistry and Biophysics, Texas A&M University, College Station, Texas, 77843, United States

**Robert L. Rider** - Department of Chemistry, Texas A&M University, College Station, Texas, 77843, United States

**Carter Lantz** - Department of Chemistry, Texas A&M University, College Station, Texas, 77843, United States

**Arthur Laganowsky** - Department of Chemistry, Texas A&M University, College Station, Texas, 77843, United States

## TOC Graphic

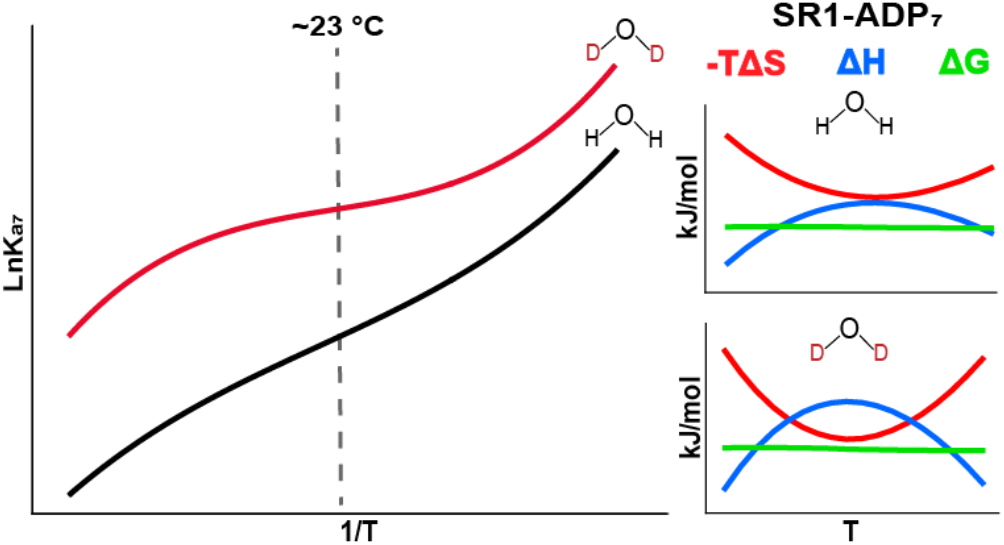

## Notes

### Competing Interest Statement

The authors have declared no competing interest.

