## Supporting Information for "Beyond Structure: Protein and Solution Dynamics Shape Ligand-Binding Thermodynamics"

### Table of Contents

| | Page |
|---|---|
| **Figure S1.** $Z_{avg}$ Vs. Temperature plots for SR1(ADP)$_n$ complexes in $H_2O$ and $D_2O$ | S2 |
| **Figure S2.** Van't Hoff plots for SR1(ADP)$_n$ complexes in $H_2O$ | S2 |
| **Figure S3.** Van't Hoff plots for SR1(ADP)$_n$ complexes in $D_2O$ | S3 |
| **Figure S4.** $\Delta C_p$ vs T plots for SR1(ADP)$_n$ complexes in $H_2O$ | S3 |
| **Figure S5.** $\Delta C_p$ vs T plots for SR1(ADP)$_n$ complexes in $D_2O$ | S4 |
| **Figure S6.** Plots of $-T\Delta S$, $\Delta H$, and $\Delta G$ vs T for SR1(ADP)$_n$ in $H_2O$ | S4 |
| **Figure S7.** Plots of $-T\Delta S$, $\Delta H$, and $\Delta G$ vs T for SR1(ADP)$_n$ in $D_2O$ | S5 |
| **Figure S8.** Intact hydrogen deuterium exchange of SR1 over time | S5 |
| **Figure S9.** Mass Spectrum of SR1 in $H_2O$ and $D_2O$ | S6 |
| **Figure S10.** Actual Vs. Input temperature for vT-ESI Device for $H_2O$ and $D_2O$ | S6 |
| **Table S1.** Deconvoluted mass of SR1 and SR1-ADP$_n$ in $H_2O$ and $D_2O$ | S7 |
| **Table S2.** Full width half maximum of the deconvoluted MS peaks of SR1 and SR1-ADP$_n$ complexes in $H_2O$ and $D_2O$ | S7 |

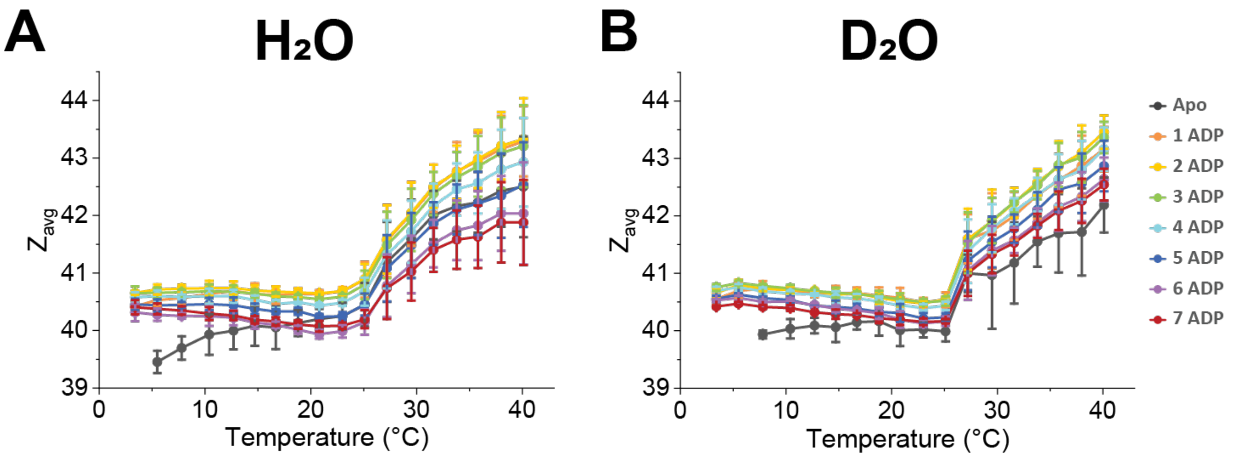


**Figure S1.** Effects of solution temperature contained in the ESI emitter on the average charge state ($Z_{avg}$) for SR1(ADP)$_n$ ($n$ = 1-7) in (**A**) $H_2O$ and (**B**) $D_2O$ with 20 μM ADP.

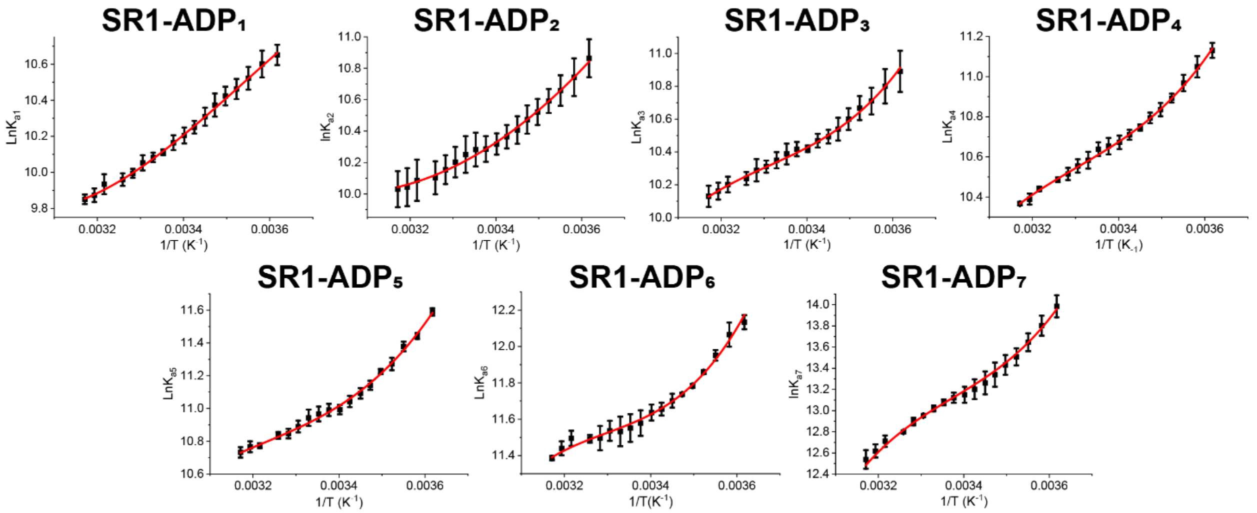


**Figure S2.** Van't Hoff plots of SR1(ADP)$_{1\text{-}7}$ binding in $H_2O$ with polynomial fits shown in red. A transition point can be observed at ~20 °C (0.0034 $K^{-1}$) in each plot.

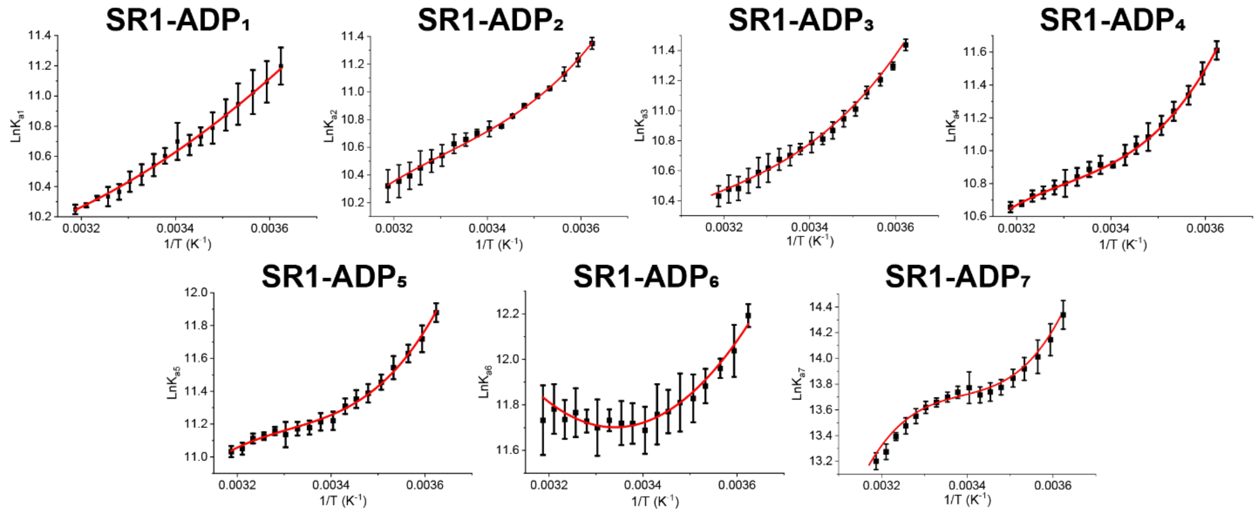


**Figure S3.** Van't Hoff plots of SR1(ADP)$_{1\text{-}7}$ binding in $D_2O$ with polynomial fits shown in red. A transition point can be observed at ~20 °C (0.0034 $K^{-1}$) in each plot.

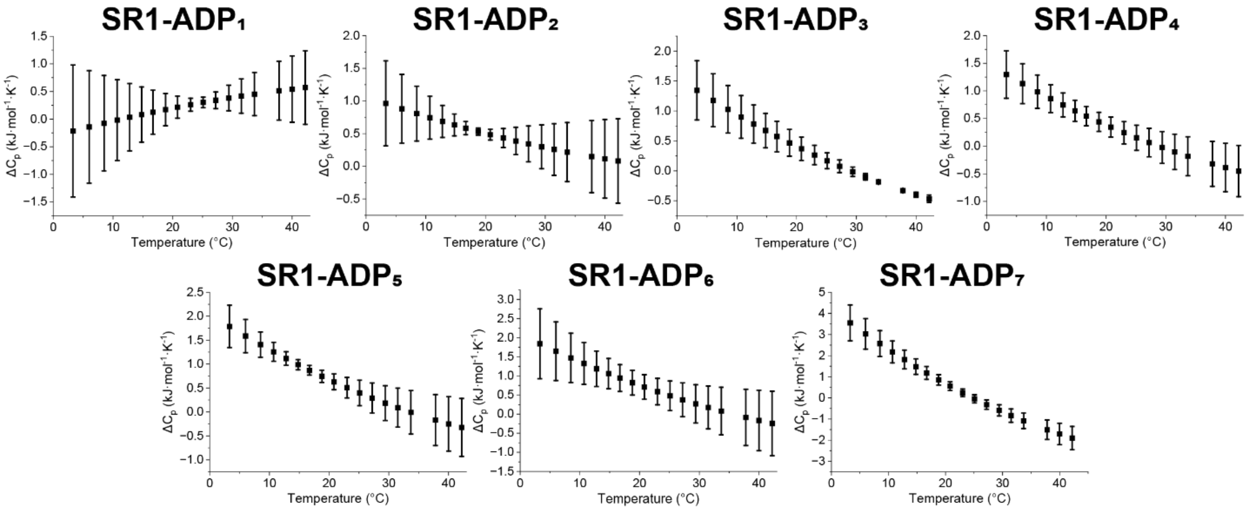


**Figure S4.** Plots showing $\Delta C_p$ in relation to temperature for SR1(ADP)$_{1\text{-}7}$ in $H_2O$ calculated from the derivative of $\Delta H$ vs T plot. The larger $\Delta C_p$ error bars observed, particularly at the temperature extremes, reflect increased sensitivity of the thermodynamic response to changes in the underlying molecular ensemble, including protein conformational dynamics and interactions with the surrounding solution environment. Furthermore, because some binding reactions exhibit relatively small changes in $\Delta C_p$, the limited numerical range of these plots visually accentuates the apparent magnitude of the error bars.

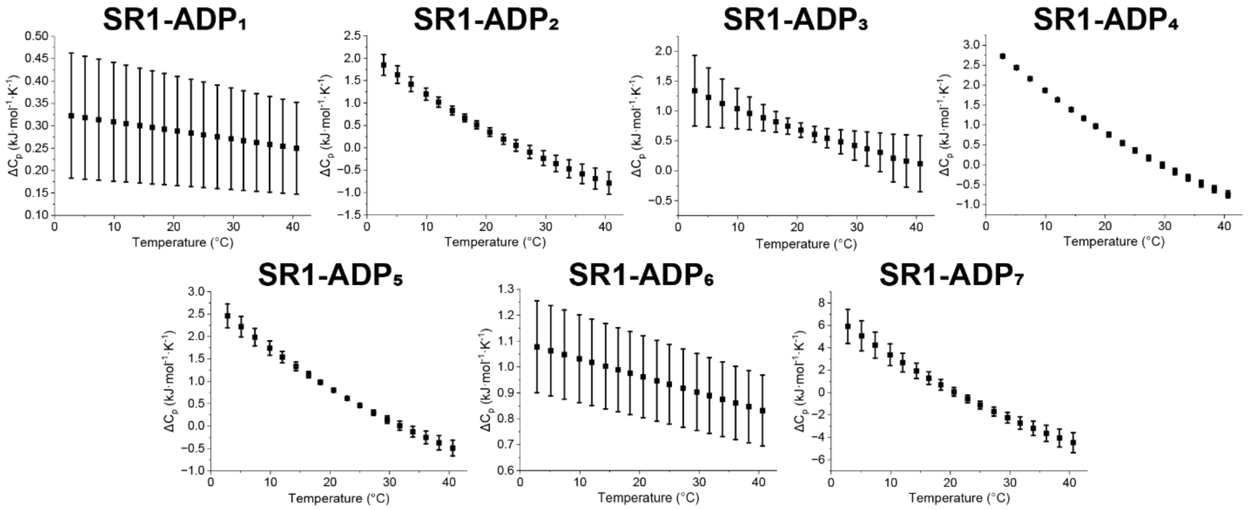


**Figure S5.** Plots showing $\Delta C_p$ in relation to temperature for SR1(ADP)$_{1\text{-}7}$ in $D_2O$ calculated from the derivative of ΔH vs T plot. The larger $\Delta C_p$ error bars observed, particularly at the temperature extremes, reflect increased sensitivity of the thermodynamic response to changes in the underlying molecular ensemble, including protein conformational dynamics and interactions with the surrounding solution environment. Furthermore, because some binding reactions exhibit relatively small changes in $\Delta C_p$, the limited numerical range of these plots visually accentuates the apparent magnitude of the error bars.

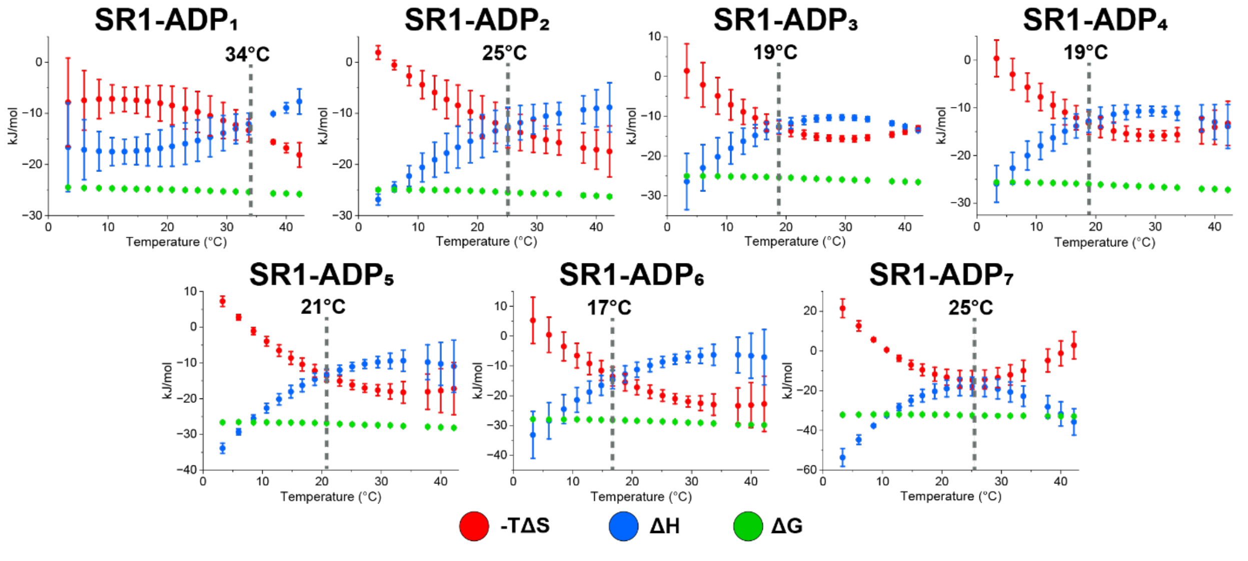


**Figure S6.** Plots showing –TΔS, ΔH, and ΔG in relation to temperature for SR1(ADP)$_{1\text{-}7}$ in $H_2O$.

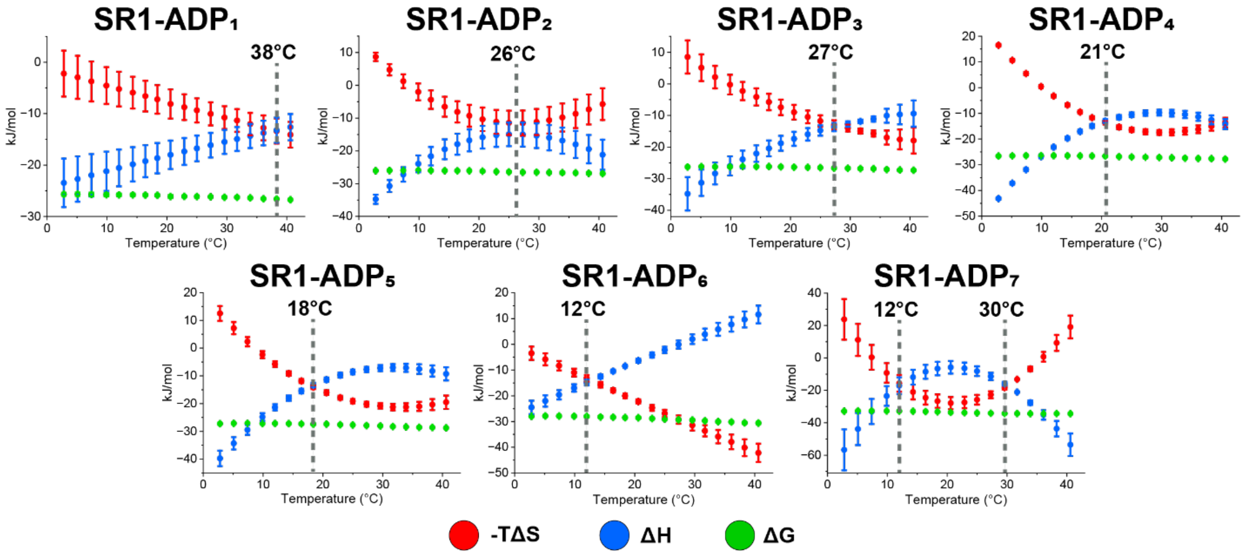


**Figure S7.** Plots showing –TΔS, ΔH, and ΔG in relation to temperature for SR1(ADP)$_{1\text{-}7}$ in $D_2O$.

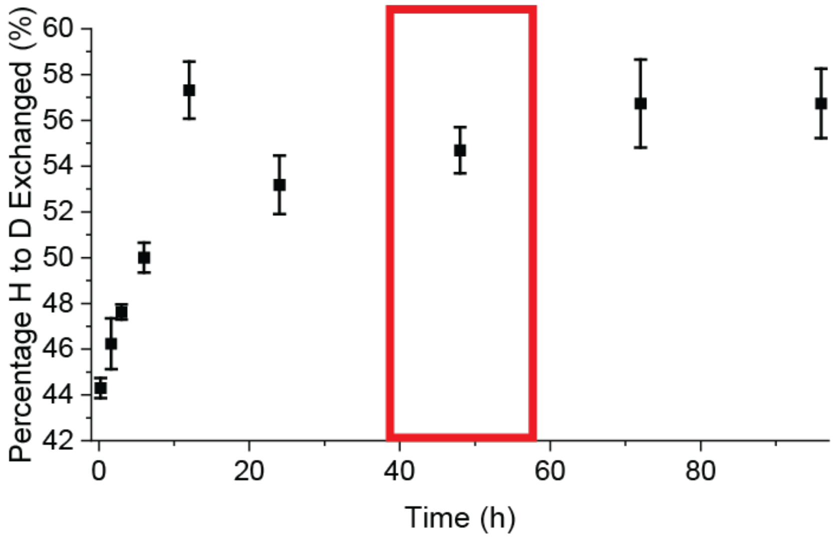


**Figure S8.** SR1 was diluted in AmAc in $D_2O$, and the percentage of exchangeable hydrogen exchanged with deuterium is plotted. The red box highlights the time frame at which SR1-ADP experiments occurred in $D_2O$ for this study.

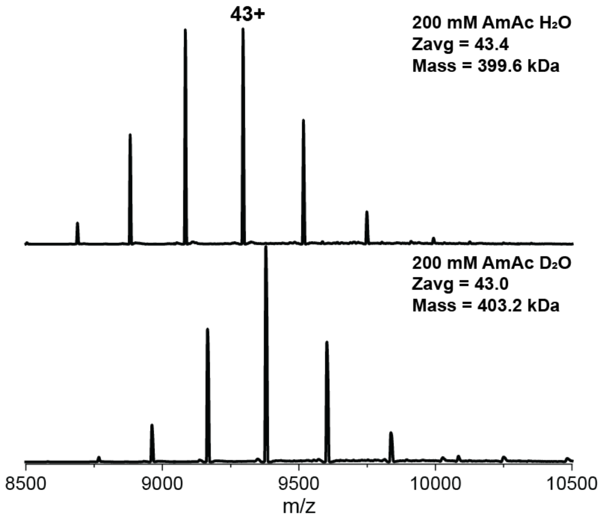


**Figure S9.** Mass spectra with mass and $Z_{avg}$ values for SR1 in $H_2O$ and $D_2O$ after being incubated at 4 °C for 48 h.

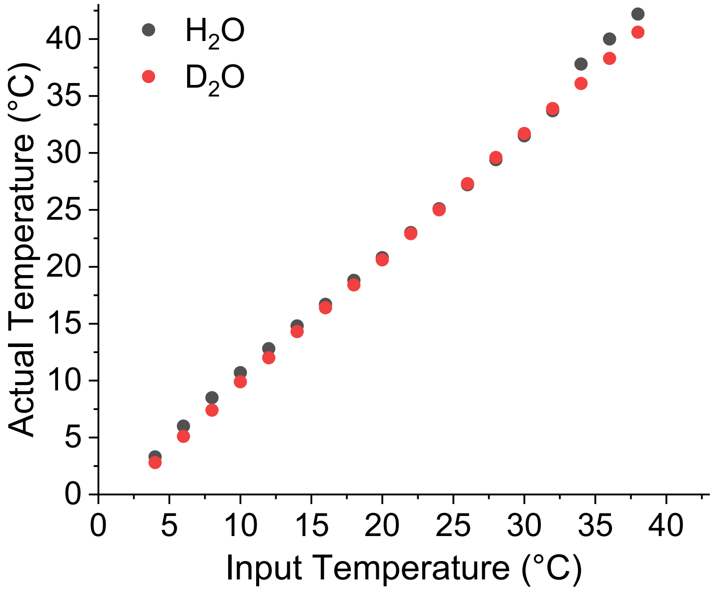


**Figure S10.** Variable-temperature electrospray ionization (vT-ESI) calibration was performed for both $H_2O$ (black) and $D_2O$ (red) analyses to account for variance between the input temperature and the actual temperature of the solution. The actual temperature was measured using a calibrated T-Type thermocouple paired to a thermocouple positioned inside the borosilicate pulled glass capillary filled with 1 uM SR1 and 1 mM $MgAc_2$ in 200 mM AmAc in $H_2O$ or $D_2O$. The actual temperatures were used for all calculations and figures.

**Table S1.** Deconvoluted mass (kDa) of SR1-ADP$_n$ in $H_2O$ and $D_2O$ after 48 h of incubation at 4 °C. ΔMass represents the change in mass of each ADP addition in relation to the reported Apo SR1 mass. The values are the averages of triplicated data sets.

| | $H_2O$ | | | 80% $D_2O$ | | |
|---|---|---|---|---|---|---|
| | Mass (kDa) | Mass Standard Deviation (Da) | ΔMass (Da) | Mass (kDa) | Mass Standard Deviation (Da) | ΔMass (Da) |
| SR1 (apo) | 399.70 | 28 | | 403.13 | 22 | |
| SR1-ADP$_1$ | 400.16 | 49 | 459 | 403.60 | 22 | 467 |
| SR1-ADP$_2$ | 400.61 | 47 | 907 | 404.06 | 29 | 928 |
| SR1-ADP$_3$ | 401.07 | 47 | 1,365 | 404.52 | 26 | 1,386 |
| SR1-ADP$_4$ | 401.53 | 58 | 1,826 | 404.98 | 25 | 1,845 |
| SR1-ADP$_5$ | 401.99 | 49 | 2,290 | 405.44 | 22 | 2,313 |
| SR1-ADP$_6$ | 402.45 | 64 | 2,749 | 405.89 | 23 | 2,761 |
| SR1-ADP$_7$ | 402.91 | 60 | 3,207 | 406.34 | 29 | 3,211 |

**Table S2.** Full width at half maximum of each peak for the deconvoluted mass spectra of SR1-ADP$_n$ in $H_2O$ and $D_2O$ (kDa).

| | $H_2O$ | 80% $D_2O$ |
|---|---|---|
| SR1 (apo) | 0.24 | 0.36 |
| SR1-ADP$_1$ | 0.22 | 0.24 |
| SR1-ADP$_2$ | 0.22 | 0.23 |
| SR1-ADP$_3$ | 0.22 | 0.23 |
| SR1-ADP$_4$ | 0.22 | 0.23 |
| SR1-ADP$_5$ | 0.23 | 0.23 |
| SR1-ADP$_6$ | 0.27 | 0.21 |
| SR1-ADP$_7$ | 0.30 | 0.29 |
